# A polyphenol-rich extract of Olive Mill Wastewater Enhances cancer chemotherapy effects, while mitigating cardiac toxicity

**DOI:** 10.1101/2021.04.08.438946

**Authors:** Adriana Albini, Marco M. G. Festa, Nadja Ring, Denisa Baci, Michael Rehman, Giovanna Finzi, Fausto Sessa, Serena Zacchigna, Antonino Bruno, Douglas M. Noonan

## Abstract

**Background:** Cardiovascular toxicities still remain one of the most undesirable side effects in cancer patients receiving chemotherapy, and cardiotoxicity has been detected associated with many therapeutic regimens. A number of mechanisms are reported for these effects, some of which are related to inflammation, oxygen radical generation, mitochondrial damage. Extra-virgin olive oil (EVOO) is rich in cancer preventive polyphenols endowed with anti-inflammatory, antioxidant activities which could exert protective effects on the heart cells. One very interesting derivative of EVOO preparation is represented by purified extract form waste waters. Here, we investigated the anti-cancer activity when combined with chemotherapeutics as well as potential cardioprotective activities of a polyphenol-rich extract from waste product of the EVOO, named A009.

**Methods and Results:** Mice bearing prostate cancer (PCa) xenografts were treated with cisplatin with and without A009. Tumor cell growth was reduced by cis and by A009 and further hindered by the combination. The effects of the A009 extract on cardiovascular toxicities was investigated in vivo. Hearts of mice were analyzed, and the mitochondria were studied by transmission electron microscopy. A protection activity by A009 was observed. To confirm the in vivo data obtained with cisplatin therapy, tumor cell lines and rat cardiomyocytes were treated with cisplatin in vitro with and without A009. A009 enhanced cisplatin and 5FU reduced cancer cell growth while did not further affect co-treated rat cardiomyocytes. Another frequently used chemotherapeutic agent 5-fluorouracil (5FU), was also tested in this assay a similar effects were observed. The cardioprotective effects of the A009 extract towards 5 FU chemotherapy were further investigated in a second system of *in vitro* cultures, on cardiomyocytes freshly isolated from mice pups. These cells were treated with 5-fluorouracil and A009. Wastewater extract mitigated toxicity of the fluorpyrimidine.

**Conclusions:** *In vivo*, we found synergisms of A009 and cisplatin in prostate cancer treatment. Hearts of mice xenografted with PCa cell lines and receiving co-treatments of A009 extracts along with cisplatin had reduced mitochondria damage compared to chemotherapy alone, indicating a cardioprotective role. A009 in vitro was additive to cisplatin and 5FU to reduce cancer cell growth while did not further affect rat cardiomyocytes cell cultures treated with cisplatin and 5FU. The A009 extract also rescued the proliferation rate of neonatal murine cardiomyocytes treated with 5-Fluorouracil. Our study demonstrates that the polyphenol rich purified A009 extracts enhances the effect of chemotherapy *in vitro* and *in vivo* but mitigates effects on heart and heart cells. It could therefore represent a potential candidate for cardiovascular prevention in patients undergoing cancer chemotherapy.

## 1. Introduction

Cancer therapy has made remarkable advances for the treatment of solid and hematological tumors, leading to significant progresses in the reduction of tumor recurrences [1–7]. Although the introduction of different antineoplastic agents in the clinic, such as monoclonal antibodies and tyrosine kinase inhibitors, has significantly augmented life expectancy [8], cardiovascular toxicities remain a major clinical concern, sometimes generating higher morbidity and mortality than tumor recurrences [8]. Cardiovascular toxicities, defined as “toxicities affecting the heart” are among the most frequent undesirable effects on cancer chemotherapy. Major effects of chemotherapy-induced cardiovascular toxicities include arrythmias, myocardial ischemia, coronary artery diseases, hypertension, and myocardial dysfunctions [7].

A major problem in the manifestation of clinically evident cardiotoxic events is the fact that they are often asymptomatic, and therefore negatively impact the cardiological prognosis of cancer patients as well as significantly limiting applicable treatment options [18,21–24]. In fact, even minor cardiac dysfunctions significantly restrict the choice of therapeutic programs, forcing the selection of those considered less aggressive and, as such, potentially less effective [1–7]. Occurrence of chemotherapy-induced cardiotoxicity is continuously increasing, as a consequence of the growing number of patients undergoing chemotherapy and the introduction of new, more aggressive anticancer drugs, often administered in combination with other toxic compounds [1–7].

This knowledge suggested that a strict dialogue between the oncologists and the cardiologists is necessary, when selecting the proper chemotherapy intervention as well as cardiac monitoring in cancer patients, bringing to a new discipline termed cardio-oncology [2].

Mitochondria represent the metabolic engine, governing and sensing the cellular energy requirements during physiological and pathological conditions [9,10]. The maintenance of mitochondrial membrane potential is crucial to supply gradients for ATP synthesis [11]. Oxidative stress, a major hallmark of age- and chronic inflammatory-related disorders and significantly impact on mitochondrial functionality [11]. Generation of ROS and mitochondrial damage are major drivers of chemotherapy-induced cardiotoxicities [12–15].

Polyphenols act as antioxidants by contrasting the generation of reactive oxygen species (ROS) [5] that drive cellular and mitochondrial damage.

It has been widely demonstrated that adherence to the Mediterranean diet is associated with a reduced risk of developing cardiovascular diseases. In recent decades, numerous epidemiological and interventional studies have confirmed this observation, underlining the close relationship between the Mediterranean diet and cardiovascular diseases [16–18]. In this context, extra virgin olive oil (EVOO), the most representative component of this diet, seems to be important in reducing the incidence of cardiovascular events, including myocardial infarction and stroke [19]. Current research on the beneficial effect of EVOO is focused on defining its protective effects against cardiovascular risk factors, such as inflammation, oxidative stress, coagulation, platelet aggregation, fibrinolysis, and endothelial or lipid dysfunction. A further approach is based on the modulation of conditions that predispose people to cardiovascular events, such as obesity, metabolic syndrome or type 2 diabetes mellitus, and chemotherapy [18,20–23]. The protective activity of EVOO results from high levels of phenolic compounds, monounsaturated fatty acids (MUFA) and other minor compounds present in EVOO [19].

Industrial EVOO processing is associated with the generation of large volume of liquid waste product, termed olive mill wastewater (OMWW) [24,25]. OMWW are rich in water soluble polyphenols endowed with anti-bacterial, anti-antioxidant, cytoprotective, [26–28] thus representing a valid waste product to be repositioned in the market.

Here, we investigate the potential cardioprotective activity of a polyphenol-rich, EVOO-derived antioxidant extract (A009), obtained from olive mill wastewater (OMWW). Extracts from A009, have been reported to exhibit chemopreventive and angiopreventive properties, *in vitro* and *in vivo*, in different cancer types [29,30]

We examine A009 effect on tumor growth when combined with a chemotherapeutic agent and evaluated the effect of the combination on the heart and cardiomyocytes, at both a cellular and molecular level, using *in vivo* (mice with prostate tumors) and *in vitro* models.

## 2. Materials and Methods

### 2.1 Chemicals

Cis-Diammine platinum dichloride (Cis-Pt) and 5-Fluorouracil (5FU), all purchase by SIGMA Aldrich were dissolved in Dimethyl sulfoxide (DMSO) and used for *in vitro* experiments as detailed below. 3-(4,5-dimethylthiazol-2-yl)-2,5-diphenyltetrazolium bromide (MTT) was purchased by SIGMA Aldrich and resuspended at 5 mg/ml. A009 polyphenol -rich extract, derived from olive mill wastewater (OMWW) processing, were purchased by Azienda Agricola fattoria La Vialla, Castiglion Fibocchi, Arezzo Italy.

### 2.2 Preparation of A009 extracts

The A009 was obtained from the OMWW derived from the processing of EVOO. Extraction procedures and polyphenol quantification has been previously published [25–27]. The polyphenol composition is not altered, following different years of cultivars [25–27]. Polyphenol content of the A009 extract is showed in supplemental table 1 and has been published [30,31].

**Table 1.** Monitoring of healthy conditions during in vivo treatments. The healthy state on mice receiving single agent (A009, dilution 1:250) alone, or Cisplatin (7 mg/Kg) alone, or the combinations of Cisplatin and the A009 extract was daily monitored. As readout of clinical parameters, the presence of skin peeling, dissertation, alterations of water and food consumption, alteration in solid (feces) and liquid (urine) dejection are showed. Data are presented as (number of events)/total animal per experimental conditions

|  | NT | Cisplatin<br>(7mg/kg) | A009 1:250 | Cisplatin (7mg/kg) +<br>A009 1:250 |
| --- | --- | --- | --- | --- |
| **Skin peeling** | 0 (10) | 5 (9) | 1 (10) | 2 (10) |
| **Dehydration** | 0 (10) | 0 (9) | 0 (10) | 0 (10) |
| **Alterations in**<br>**water**<br>**consumption** | 0 (10) | 0 (9) | 0 (10) | 0 (10) |
| **Alterations in food**<br>**consumption** | 0 (10) | 0 (9) | 0 (10) | 0 (10) |
| **Urine** | 0 (10) | 0 (9) | 0 (10) | 0 (10) |
| **Feces** | 0 (10) | 0 (9) | 0 (10) | 0 (10) |

### 2.3 Cell line culture and maintenances

The human prostate cancer (PCa) cell lines DU-145, 22Rv1 and the colorectal cancer cell line HT29 (all purchased by ATCC) were maintained in RPMI 1640 medium, supplemented with 10% Fetal Bovine Serum (FBS) (Euroclone), 2 mM l-glutamine (Euroclone), 100 U/ml penicillin and 100 µg/ml streptomycin (Euroclone), at 37°C, 5% CO2. The rat cardiomyocyte cell line H9C2 (PromoCell) was maintained in Myocyte Growth Medium medium plus Myocyte supplements mix (PromoCell), addition with 10% Fetal Bovine Serum (FBS) (Euroclone), 2 mM L-glutamine (Euroclone), 100 U/ml penicillin and 100 μg/ml streptomycin (Euroclone), at 37°C, 5% CO_2_. Cells were routinely screened for eventual mycoplasma contaminations.

### 2.4 Detection of cardioprotective activities in in vivo tumor xenograft models

We used a mouse model of prostate cancer to determine whether co-treatment with the chemotherapeutic agent cisplatin and A009 extract could exert a protective effect on the hearts of the treated animals. The effects of the A009 extracts in inhibiting prostate cancer (PCa) tumor cell growth was assessed using an *in vivo* xenograft model. 5-week-old male Nu/MRI nude mice (from Charles River) were used, with four animals per experimental group. Animals were housed in a conventional animal facility with 12:12 h light dark cycles and fed ad libitum. Animals were subcutaneously injected into the right flank with 2.5 × 10^6^ 22Rv1 cells or DU-145 cells, in a total volume of 300 μl, containing 50% serum free RMPI 1650, and 50% 10 mg/mL reduced growth factor Matrigel (Corning) with or without A009 (dilution 1:250). From day 0 animals received A009 daily (dilution 1:250), in the drinking water. When tumors were palpable, mice received Cisplatin, 7 mg/kg *i.p*, twice a week. At day 27, the tumor cell growth was stopped, tumors were excised, weighted and tumor volume was measured with a caliper and determined using the formula (W^2^ × L)/2. Hearts were surgically removed from animals and used for transmission electron microscopy analyses.

All the procedures involving the animals and their care were performed according to the institutional guidelines, in compliance with national and international law and guidelines for the use of animals in biomedical research and housed in pathogen-free conditions. All the procedures applied were approved by the local animal experimentation ethics committee (ID# #06_16 Noonan) of the University of Insubria and by the Italian Health Ministry (ID#225/2017-PR).

### 2.5 Transmission Electron Microscopy analysis of murine hearts

Hearts were surgically excised from animals and extensively washed in PBS. Heart sections were obtained using a scalpel and then placed in fixing solution for TEM processing (2% PFA, 2% glutaraldehyde), finally post-fixed using 1% osmium tetroxide and embedded in an Epon-Araldite resin. Following exposure to uranyl acetate and lead citrate, thin sections were analyzed by TEM, using a Morgagni electron microscope (Philips) at 3500X magnification, to detect mitochondrial alterations in terms of morphology, size, organization and quantity. The number of altered mitochondria per section, exhibiting altered morphology/shape, was counted using the ImageJ software.

### 2.6 Combination effect of chemotherapy and A009 on cancer cell lines

To investigate whether the A009 extract could synergize with chemotherapy, the prostate cancer DU-145 cell line or the colorectal cancer HT-29 cell line were treated with Cis-Pt 100 μM or 5-FU 100 μM, respectively, alone or in combination with A009 L3 or L4 extracts, for 24 to 72 hours. Detection of cell viability was determined by MTT (3-[4,5-dimethylthiazole-2-yl]-2,5-diphenyltetrazolium bromide) assay, on 3,000 cardiomyocytes/well, seeded into a 96 well plate.

### 2.7 Effects of A009 extracts on adult rat cardiomyocyte

To evaluate the effects of the A009 extracts on chemotherapy induced cardiotoxicity, after preliminary experiment to assess dosages, adult rat cardiomyocyte H9C2 cells were treated with 5-FU 100 μM or Cis-Pt 100 μM, alone or in combination with A009 L3 or L4 extracts, for 24 to 72 hours. The schedule treatments included a prevention approach by pre-treating cardiomyocyte with A009 L3 and L4 extracts at T24 to T48 h, subsequently A009 L3 or L4 extracts were removed, and wells were auditioned with fresh medium containing Cis-Pt 100 μM or 5-Fu100 μM. Detection of cell viability was determined by MTT assay descripted in 2.6.

### 2.8 Isolation of neonatal murine cardiomyocytes

Cardiomyocytes were isolated from neonatal C57/Bl6 mice at 2 days after birth as previously described, with minor modifications [32]. Briefly, hearts were removed and cleaned in calcium and bicarbonate-free Hanks’ balanced salt solution with Hepes (CBFHH, containing 137 mM NaCl, 5.36 mM KCl, 0.81 mM MgSO_4_ 7H_2_O, 5.55 mM dextrose, 0.44 mM KH_2_PO_4_, 0.34 mM Na_2_HPO_4_ 7H_2_O, and 20.06 mM HEPES). Excess blood and valves were removed, and hearts were diced. The tissue was then enzymatically digested using CBFHH supplemented with 1.75 mg/ml of Trypsin (BD Biosciences) and 20 mg/ml of DNAse I (Sigma). Tissue was digested for 3 hours, with cells harvested into fetal bovine serum (FBS) every 10 min to stop the digestion. Cells were then filtered using a 40 μm cell strainer and pre-plated for 2 h to remove contaminating fibroblasts. Finally, cardiomyocytes were collected and seeded on tissue culture plates treated for primary cultures. Cells were cultured in Dulbecco’s modified Eagle medium 4.5 g/L glucose (DMEM, Life Technologies) supplemented with 5% FBS, 20 mg/ml vitamin B12 (Sigma), 100 U/ml penicillin and 100 mg/ml streptomycin (Sigma).

### 2.9 Effects of A009 extracts on neonatal murine-derived cardiomyocytes

To evaluate the effect of the A009 extract on cardiomyocyte viability *in vitro*, 30,000 cardiomyocytes/well were seeded into a 96 well plate. One day after plating, cells were treated with L3 and L4 A009 extracts, dilution of 1:800, for 24 h. On day 2, cells were treated with 4.6 μM of 5-Fluorouracil. Following 24 and 48 hours, cells were fixed and stained using anti-Cardiac Troponin I antibody (Abcam, ab47003, dilution of 1:200) and Hoechst 33342 (Invitrogen, H3570, dilution of 1:5000). The number of cardiomyocytes for each time point was counted in three independent experiments.

## 3. Results

### 3.1 Cardioprotective activities of A009 extracts in in vivo models of cardiotoxicity induced by anticancer drug

We used a mouse model of prostate cancer to determine the A009 effect on tumors and the hearts of mice treated with the chemotherapeutic agent cisplatin. During the treatment schedule, we did not observe behavioral changes, alterations in food intake, water consumption, or dejections by the animals included in all the experimental groups of the study (Table 1).

Animals receiving the different treatment did not show weight loss during the tumor cell growth kinetic (Figure 1A). Interesting, A009 also reduced the skin peeling induced by cisplatin treatments (from 5/9 mice to (2/10 mice) (Table 1). We found that the combination of cisplatin with the A009 extract synergized further reduced the PCa cell tumor weight, as compared to the treatment with cisplatin alone (Figure 1 B). The morphological analysis did not reveal macroscopic differences amongst the hearts of the various experimental groups (data not shown).

**Figure 1:**
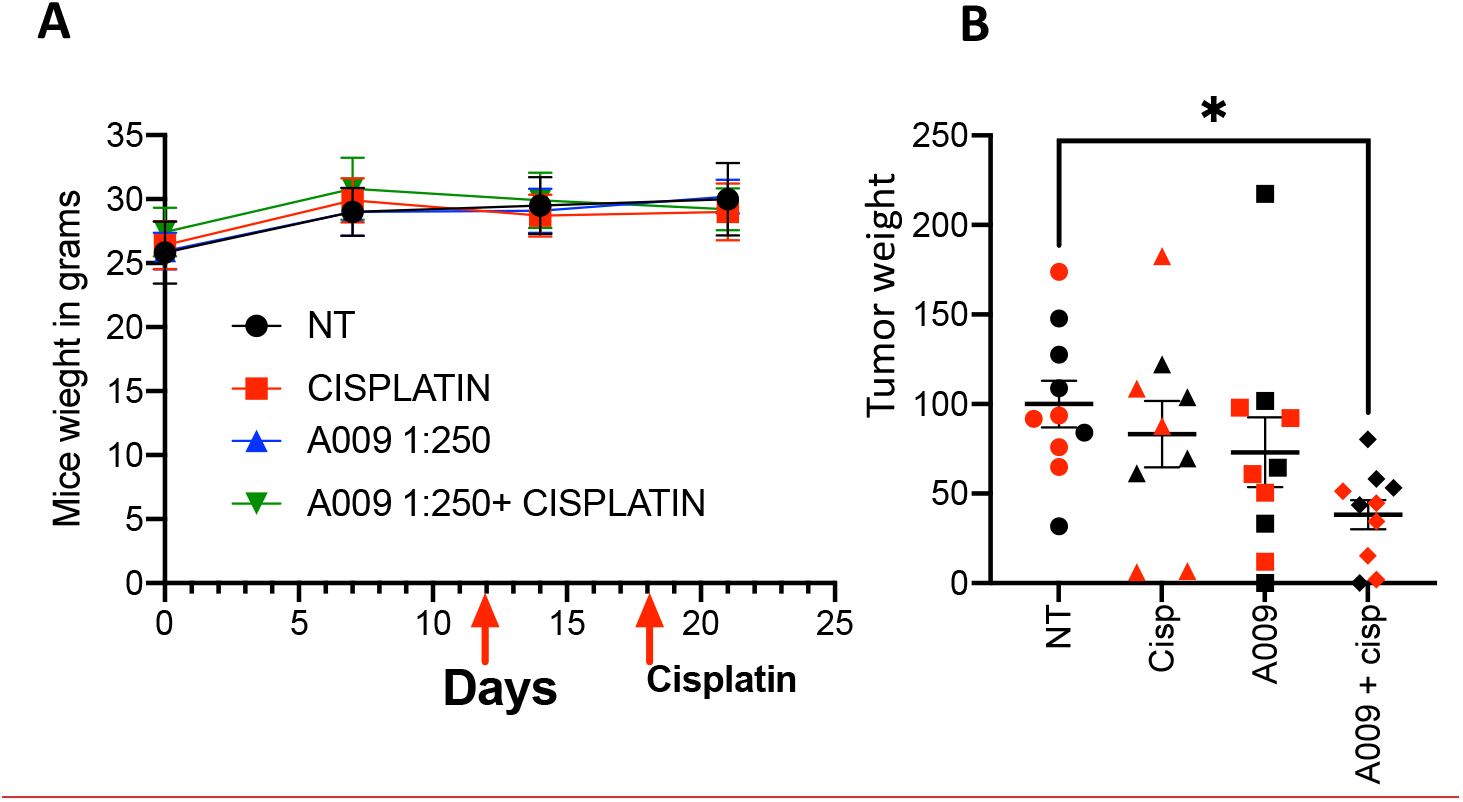
Dietary administration of the A009 extract, in combination with chemotherapy, result in both synergism by reducing tumor weight. Dietary administration (drinking water) of A009 extracts synergize with chemotherapy by reducing tumor weight *in vivo*. In panel A the red arrows indicate the dose of the cisplatin (7 mg/Kg), the mice weights were same. In panel B, the effects of the combination of A009 extract with cisplatin (7 mg/Kg), was determined using an orthotopic *in vivo* model of prostate cancer cells DU-145 (red), 22Rv1 (black). Data are showed as mean ± SEM, one-way ANOVA, *p<0.05.

However ultrastructural analysis, using transmission electron microscopy (TEM), showed that animals treated with cisplatin which received also the A009 extract have a reduced number of damaged mitochondria (showing a rounder shape and having mitochondrial cristae better organized and higher in number), as compared to the hearts of mice treated with cisplatin only (Figure 2A-B). We also observed a more regular muscle fiber disposition in the hearts of animals treated with A009 and cisplatin as compared to those treated with cisplatin alone.

**Figure 2:**
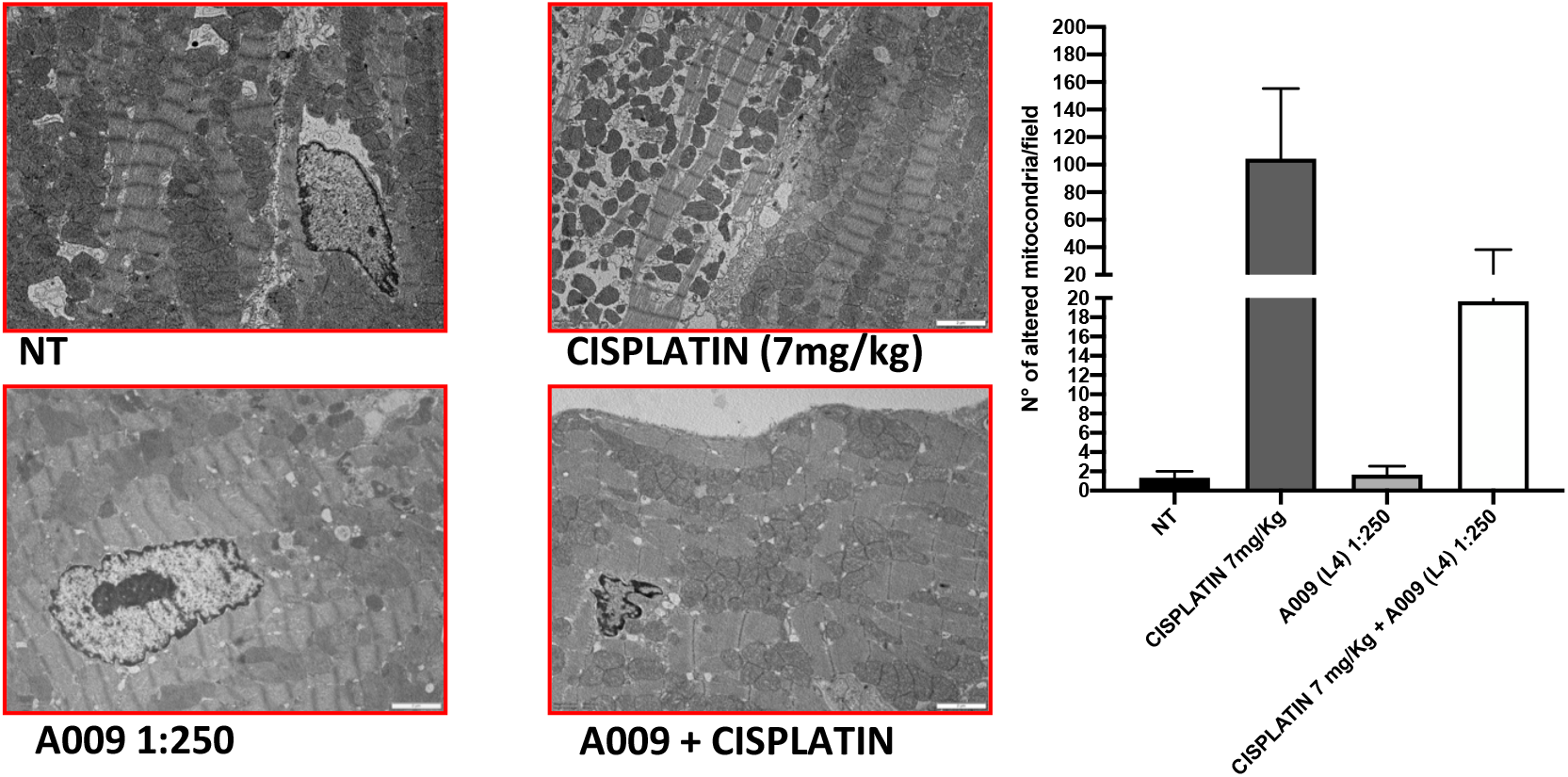
A009 cardioprotective activities against cisplatin-induce cardiotoxicity *in vivo*. Mitochondria number, shape/morphology and color was monitored, by transmission electron microscopy (TEM) on hearts from mice treated with cisplatin alone (7 mg/Kg), A009 extract (dilution 1:250, in drinking water) or the cisplatin-A009 extract combination. Data are showed as mean ± SEM, one-way ANOVA, *p<0.05, **p<0.01, ***p<0.001, ****p<0.0001. A009 batch extract; NT: vehicle control.

### 3.2 A009 activities against tumor cell lines and heart cell lines

Cisplatin and 5-FU treatment decreased both prostate and colon cancer cell growth; The proliferation of the tumors treated with A009 was also significantly different from the vehicle control. A009 enhanced the effect of the cisplatin and 5-FU alone (Figure 3). 5-FU and cisplatin were toxic to rat cardiomyocytes, while the A009 was not. Furthermore, A009 added to Cisplatin or 5FU did not enhance the growth reduction (Figure 3). Importantly, while A009 significantly diminishes tumor cell proliferation rate and has additive effect with cisplatin and 5-FU, it does not affect myocyte growth and it does not enhance toxicity of cisplatin and 5-FU (Figure 3).

**Figure 3:**
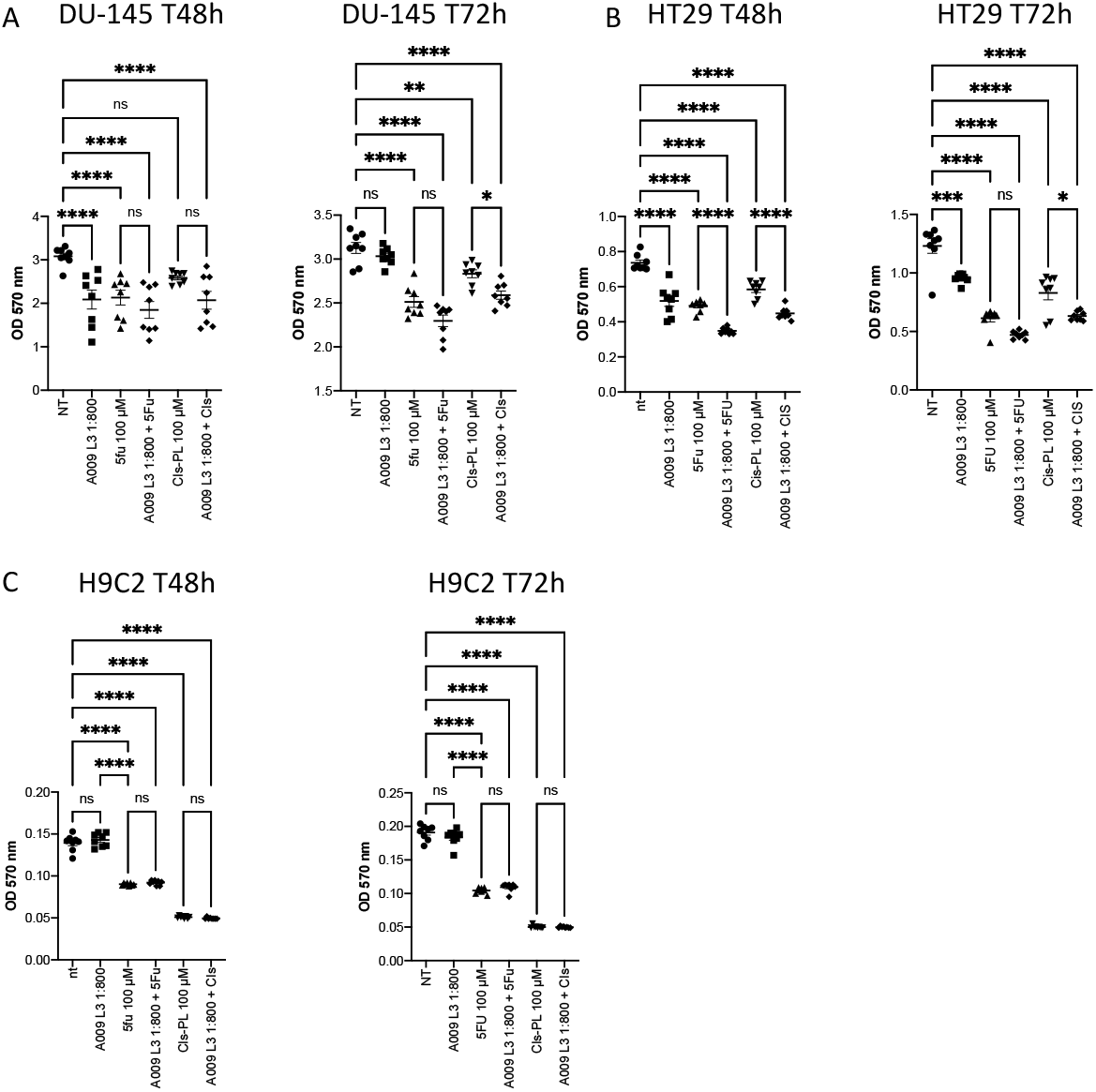
Activities of A009 extracts combined with chemotherapy on tumor cells and cardiomyocytes. A009 (batch L3) decease the proliferation rate of tumors cells *in vitro* (A: DU-145 PCa and B: HT29 CRC) and has additive effects on the cisplatin and the 5-Fluorouracil (5FU) effects. The cardiomyocytes proliferation rate is not affected by A009 alone (C), and reduced proliferation by 5FU and cisplatin is not enhanced by A009 *in vitro*. Data are showed as mean ± SEM, one-way ANOVA, *p<0.05, **p<0.01, ***p<0.001, ****p<0.0001.

### 3.3 Protective activities of A009 extracts on neonatal murine cardiomyocytes

We observed a cardioprotective effect of the A009 extract in neonatal murine cardiomyocytes, following co-treatment with the chemotherapeutic drugs 5-fluorouracil (5-FU). The protective effect of the A009 extract was determined by quantifying the number of surviving cardiomyocytes at 24 hours (Figure 2 A) and 48 hours (Figure 2 B) post treatment. At the early time point of 24 h, A009 showed a cardioprotective effect in basal conditions, and was slightly protective against 5-FU (Figure 4). After 48 h, A009 was consistently cardioprotective against both 5-FU (Figure 4)

**Figure 4:**
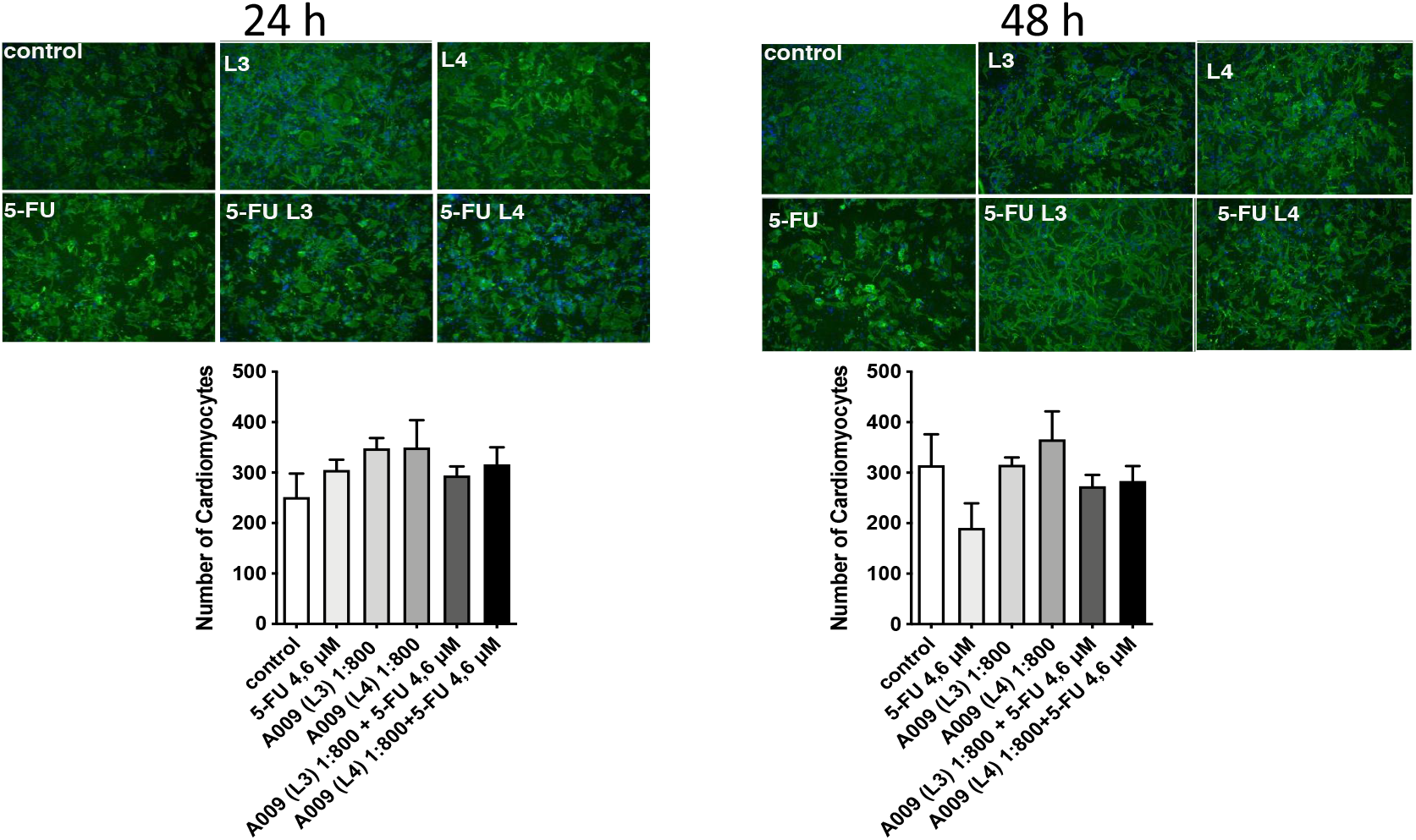
Protective activities of A009 extracts on neonatal murine cardiomyocytes. The cardioprotective effects of the A009 extract on chemotherapy induced cardiotoxicities was assessed, *in vitro*, on neonatal murine cardiomyocytes. Neonatal murine cardiomyocytes were exposed for 24 hours (A) and 48 hours (B) to 5-FU (4,6 μM) alone, A009 extracts (dilution 1:800, batches L3 or L4) alone, or the combination of the 5-FU and A009 extracts (dilution 1:800, batches L3 or L4). Data are showed as mean ± SEM. 5-FU: 5-fluoro-Uracile; L3/L4: A009 batch extract; NT: vehicle control.

## 4. Discussion

Cardiovascular toxicities still remain a major challenge in clinical oncology [1–4,8]. While chemotherapeutic agents efficiently target malignantly transformed cells, they simultaneously induce cell death of healthy cells [1–4,8]. The cardiovascular system is the major off target of anti-neoplastic drugs [1–4,8]. Most of the cytotoxic activities of chemotherapeutic agents on normal cells are due to the induction of exacerbated oxidative stress, through the generation of both ROS and reactive nitrogen species (RNS) [3,32]. Agents such as anti-inflammatory, antioxidants, able to counteract these effects, can be used to reduce side effects from chemotherapeutics and can be easily tolerated by oncologic patients and administered by dietary regimen [33,34].

Many dietary polyphenols demonstrate antioxidant and cytoprotective properties [35–38]. We tested the ability of a polyphenol-rich purified extract of OMWW, termed A009, to protect from cardiovascular damages induced by anti-cancer agents.

Mimicking a scenario closer to the clinic, we tested the cardioprotective properties of the A009 extract in *in vivo* murine models of tumor xenograft treated with cisplatin, a chemotherapy agent associated with cardiotoxicity and mitotoxicity [39–42]. Mice subcutaneously injected with prostate cancer cell line, co-treated with A009 and the chemotherapeutic drug cisplatin showed a reduced number of abnormal and damaged mitochondria, as compared to those treated with cisplatin alone. Mitochondria have an essential role in myocardial tissue homeostasis [43,44] and diverse chemical compounds and chemotherapy drugs have been known to directly or indirectly modulate cardiac mitochondrial function [45,46]. Mitochondrial oxidative stress and dysfunction is a common mechanism in cardiotoxic effects [13,14,39,41–43].

Cisplatin was tested in vitro on prostate cancer cells, with or without A009 and its effects compared to the ones on rat cardiomyocytes.

5-FU is also a common cancer chemotherapeutic. We had studied human cardiac myocytes *in vitro* senescent phenotype and autophagic features upon 5FU treatment [6]. While A009 significantly diminished tumor cell proliferation rate and had additive effect with cisplatin and 5-FU, it did not affect myocyte growth as single treatment and it did not enhance toxicity of cisplatin and 5-FU.

Based on these results, we investigated the effects of the A009 extracts also on fresh cardiomyocytes isolated from neonatal mice. In these experiments, we validate 5-FU on cardiotoxicity [2,3,6,37,38]. We observed that cardiomyocytes co-treated with the A009 extracts and a chemotherapeutic drug 5-FU exhibited less reduction of the number of cardiomyocytes, as compared with the drug alone. This rescue was maintained from 24 to 48 hours of cardiomyocyte culture and treatment and potentially related to the antioxidant polyphenols present in the A009 extracts.

## 5. Conclusions

### 5.1 A009 extracts enhances chemotherapy effects on tumor cells in vivo and in vitro

Here, we demonstrated that the A009 extracts, although additive in cancer therapy do not have cardiotoxic effects, and actually can mitigate chemotherapy-induced cardiotoxicity. One of the effects, detected by transmission electron microscopy on hearts of the treated mice, suggests mitochondrial protection and anti-oxidant capabilities of A009.

Our study demonstrates that the polyphenol rich purified A009 extracts are a valid candidate for combination chemotherapy and for cardiovascular protection from induced cardiac damage.

---

**Supplementary Figure 1: Phenolic composition of A009 was obtained by HPLC-DADMS-MS.** Samples were analyzed by HPLC with UV–vis and MS detection. The identification of phenolic compounds from samples was carried out as previously reported by interpreting their mass spectra determined via LC-MS-MS and comparing to data reported in literature identified the compounds.

**Supplementary Figure 2: Activities of a second A009 batch on DU-145 PCa tumor cell line.** A009 (batch L4) decease the proliferation rate of DU-145 PCa tumors line cells *in vitro* and has additive effects on the cisplatin). Data are showed as mean ± SEM, one-way ANOVA, **p<0.01, ***p<0.001, ****p<0.0001.

## Author Contributions

Conceptualization, A.A, A.B, S.Z., F.S. and D.M.N; methodology, N.R., M.R., M.M. G.F, B.D., G.F.; data analysis, A.B., D.B., N.R., S.Z., M.R., DMN; resources, A.A.; data curation, AB, DB, NR, MR, M. M. G. F, DB; writing—original draft preparation, A.A., A.B., DMN; supervision, A.A., A.B., D. M. N.; funding acquisition, S. Z., A.A. All authors have read and agreed to the published version of the manuscript

## Acknowledgments

We thank Paola Corradino for support in literature research and text editing.

## References

1. Albini, A.; Donatelli, F.; Focaccetti, C.; D’Elios, M.M.; Noonan, D.M. Renal dysfunction and increased risk of cardiotoxicity with trastuzumab therapy: a new challenge in cardio-oncology. Intern Emerg Med 2012, 7, 399–401, doi:10.1007/s11739-012-0845-2.

2. Albini, A.; Pennesi, G.; Donatelli, F.; Cammarota, R.; De Flora, S.; Noonan, D.M. Cardiotoxicity of anticancer drugs: the need for cardio-oncology and cardio-oncological prevention. J Natl Cancer Inst 2010, 102, 14–25, doi:10.1093/jnci/djp440.

3. Angsutararux, P.; Luanpitpong, S.; Issaragrisil, S. Chemotherapy-Induced Cardiotoxicity: Overview of the Roles of Oxidative Stress. Oxid Med Cell Longev 2015, 2015, 795602, doi:10.1155/2015/795602.

4. Conway, A.; McCarthy, A.L.; Lawrence, P.; Clark, R.A. The prevention, detection and management of cancer treatment-induced cardiotoxicity: a meta-review. BMC Cancer 2015, 15, 366, doi:10.1186/s12885-015-1407-6.

5. Curigliano, G.; Cardinale, D.; Dent, S.; Criscitiello, C.; Aseyev, O.; Lenihan, D.; Cipolla, C.M. Cardiotoxicity of anticancer treatments: Epidemiology, detection, and management. CA Cancer J Clin 2016, 66, 309–325, doi:10.3322/caac.21341.

6. Focaccetti, C.; Bruno, A.; Magnani, E.; Bartolini, D.; Principi, E.; Dallaglio, K.; Bucci, E.O.; Finzi, G.; Sessa, F.; Noonan, D.M., et al. Effects of 5-fluorouracil on morphology, cell cycle, proliferation, apoptosis, autophagy and ROS production in endothelial cells and cardiomyocytes. PLoS One 2015, 10, e0115686, doi:10.1371/journal.pone.0115686.

7. Polonsky, T.S.; DeCara, J.M. Risk factors for chemotherapy-related cardiac toxicity. Curr Opin Cardiol 2019, 34, 283–288, doi:10.1097/HCO.0000000000000619.

8. Senkus, E.; Jassem, J. Cardiovascular effects of systemic cancer treatment. Cancer treatment reviews 2011, 37, 300–311, doi:10.1016/j.ctrv.2010.11.001.

9. Missiroli, S.; Genovese, I.; Perrone, M.; Vezzani, B.; Vitto, V.A.M.; Giorgi, C. The Role of Mitochondria in Inflammation: From Cancer to Neurodegenerative Disorders. J Clin Med 2020, 9, doi:10.3390/jcm9030740.

10. Vringer, E.; Tait, S.W.G. Mitochondria and Inflammation: Cell Death Heats Up. Front Cell Dev Biol 2019, 7, 100, doi:10.3389/fcell.2019.00100.

11. Bhatti, J.S.; Bhatti, G.K.; Reddy, P.H. Mitochondrial dysfunction and oxidative stress in metabolic disorders - A step towards mitochondria based therapeutic strategies. Biochim Biophys Acta Mol Basis Dis 2017, 1863, 1066–1077, doi:10.1016/j.bbadis.2016.11.010.

12. Hahn, V.S.; Lenihan, D.J.; Ky, B. Cancer therapy-induced cardiotoxicity: basic mechanisms and potential cardioprotective therapies. J Am Heart Assoc 2014, 3, e000665, doi:10.1161/JAHA.113.000665.

13. Ichikawa, Y.; Ghanefar, M.; Bayeva, M.; Wu, R.; Khechaduri, A.; Naga Prasad, S.V.; Mutharasan, R.K.; Naik, T.J.; Ardehali, H. Cardiotoxicity of doxorubicin is mediated through mitochondrial iron accumulation. J Clin Invest 2014, 124, 617–630, doi:10.1172/JCI72931.

14. Nitiss, K.C.; Nitiss, J.L. Twisting and ironing: doxorubicin cardiotoxicity by mitochondrial DNA damage. Clin Cancer Res 2014, 20, 4737–4739, doi:10.1158/1078-0432.CCR-14-0821.

15. Zhang, X.; Hu, C.; Kong, C.Y.; Song, P.; Wu, H.M.; Xu, S.C.; Yuan, Y.P.; Deng, W.; Ma, Z.G.; Tang, Q.Z. FNDC5 alleviates oxidative stress and cardiomyocyte apoptosis in doxorubicin-induced cardiotoxicity via activating AKT. Cell Death Differ 2020, 27, 540–555, doi:10.1038/s41418-019-0372-z.

16. Billingsley, H.E.; Carbone, S. The antioxidant potential of the Mediterranean diet in patients at high cardiovascular risk: an in-depth review of the PREDIMED. Nutr Diabetes 2018, 8, 13, doi:10.1038/s41387-018-0025-1.

17. Grosso, G.; Mistretta, A.; Frigiola, A.; Gruttadauria, S.; Biondi, A.; Basile, F.; Vitaglione, P.; D’Orazio, N.; Galvano, F. Mediterranean diet and cardiovascular risk factors: a systematic review. Crit Rev Food Sci Nutr 2014, 54, 593–610, doi:10.1080/10408398.2011.596955.

18. Estruch, R.; Ros, E.; Salas-Salvado, J.; Covas, M.I.; Corella, D.; Aros, F.; Gomez-Gracia, E.; Ruiz-Gutierrez, V.; Fiol, M.; Lapetra, J., et al. Primary Prevention of Cardiovascular Disease with a Mediterranean Diet Supplemented with Extra-Virgin Olive Oil or Nuts. N Engl J Med 2018, 378, e34, doi:10.1056/NEJMoa1800389.

19. Nocella, C.; Cammisotto, V.; Fianchini, L.; D’Amico, A.; Novo, M.; Castellani, V.; Stefanini, L.; Violi, F.; Carnevale, R. Extra Virgin Olive Oil and Cardiovascular Diseases: Benefits for Human Health. Endocrine, metabolic & immune disorders drug targets 2018, 18, 4–13, doi:10.2174/1871530317666171114121533.

20. Marcelino, G.; Hiane, P.A.; Freitas, K.C.; Santana, L.F.; Pott, A.; Donadon, J.R.; Guimaraes, R.C.A. Effects of Olive Oil and Its Minor Components on Cardiovascular Diseases, Inflammation, and Gut Microbiota. Nutrients 2019, 11, doi:10.3390/nu11081826.

21. Martinez-Gonzalez, M.A.; Gea, A.; Ruiz-Canela, M. The Mediterranean Diet and Cardiovascular Health. Circ Res 2019, 124, 779–798, doi:10.1161/CIRCRESAHA.118.313348.

22. Mazzocchi, A.; Leone, L.; Agostoni, C.; Pali-Scholl, I. The Secrets of the Mediterranean Diet. Does [Only] Olive Oil Matter? Nutrients 2019, 11, doi:10.3390/nu11122941.

23. Nediani, C.; Ruzzolini, J.; Romani, A.; Calorini, L. Oleuropein, a Bioactive Compound from Olea europaea L., as a Potential Preventive and Therapeutic Agent in Non-Communicable Diseases. Antioxidants (Basel) 2019, 8, doi:10.3390/antiox8120578.

24. El-Abbassi, A.; Kiai, H.; Hafidi, A. Phenolic profile and antioxidant activities of olive mill wastewater. Food Chem 2012, 132, 406–412, doi:10.1016/j.foodchem.2011.11.013.

25. Vougogiannopoulou, K.; Angelopoulou, M.T.; Pratsinis, H.; Grougnet, R.; Halabalaki, M.; Kletsas, D.; Deguin, B.; Skaltsounis, L.A. Chemical and Biological Investigation of Olive Mill Waste Water - OMWW Secoiridoid Lactones. Planta Med 2015, 81, 1205–1212, doi:10.1055/s-0035-1546243.

26. Belaqziz, M.; Tan, S.P.; El-Abbassi, A.; Kiai, H.; Hafidi, A.; O’Donovan, O.; McLoughlin, P. Assessment of the antioxidant and antibacterial activities of different olive processing wastewaters. PLoS One 2017, 12, e0182622, doi:10.1371/journal.pone.0182622.

27. Schaffer, S.; Muller, W.E.; Eckert, G.P. Cytoprotective effects of olive mill wastewater extract and its main constituent hydroxytyrosol in PC12 cells. Pharmacol Res 2010, 62, 322–327, doi:10.1016/j.phrs.2010.06.004.

28. Abu-Lafi, S.; Al-Natsheh, M.S.; Yaghmoor, R.; Al-Rimawi, F. Enrichment of Phenolic Compounds from Olive Mill Wastewater and In Vitro Evaluation of Their Antimicrobial Activities. Evid Based Complement Alternat Med 2017, 2017, 3706915, doi:10.1155/2017/3706915.

29. Gallazzi, M.; Festa, M.; Corradino, P.; Sansone, C.; Albini, A.; Noonan, D.M. An Extract of Olive Mill Wastewater Downregulates Growth, Adhesion and Invasion Pathways in Lung Cancer Cells: Involvement of CXCR4. Nutrients 2020, 12, doi:10.3390/nu12040903.

30. Baci, D.; Gallazzi, M.; Cascini, C.; Tramacere, M.; De Stefano, D.; Bruno, A.; Noonan, D.M.; Albini, A. Downregulation of Pro-Inflammatory and Pro-Angiogenic Pathways in Prostate Cancer Cells by a Polyphenol-Rich Extract from Olive Mill Wastewater. Int J Mol Sci 2019, 20, doi:10.3390/ijms20020307.

31. Bassani B. R.T., De Stefano D, Pizzichini D, Corradino P, Macrì N, Noonan DM, Albini A, Bruno A.. Potential chemopreventive activities of a polyphenol rich purified extract from olive mill wastewater on colon cancer cells. . Journal of Functional Foods 2016, 27, 236–248, doi:10.1016/j.jff.2016.09.009.

32. Zhang, J.; Lei, W.; Chen, X.; Wang, S.; Qian, W. Oxidative stress response induced by chemotherapy in leukemia treatment. Mol Clin Oncol 2018, 8, 391–399, doi:10.3892/mco.2018.1549.

33. Kaiserova, H.; Simunek, T.; van der Vijgh, W.J.; Bast, A.; Kvasnickova, E. Flavonoids as protectors against doxorubicin cardiotoxicity: role of iron chelation, antioxidant activity and inhibition of carbonyl reductase. Biochim Biophys Acta 2007, 1772, 1065–1074, doi:10.1016/j.bbadis.2007.05.002.

34. Vincent, D.T.; Ibrahim, Y.F.; Espey, M.G.; Suzuki, Y.J. The role of antioxidants in the era of cardiooncology. Cancer Chemother Pharmacol 2013, 72, 1157–1168, doi:10.1007/s00280-013-2260-4.

35. Baranowska, M.; Bartoszek, A. Antioxidant and antimicrobial properties of bioactive phytochemicals from cranberry. Postepy Hig Med Dosw (Online) 2016, 70, 1460–1468, doi:10.5604/17322693.1227896.

36. Krajka-Kuzniak, V.; Szaefer, H.; Ignatowicz, E.; Adamska, T.; Oszmianski, J.; Baer-Dubowska, W. Effect of Chokeberry (Aronia melanocarpa) juice on the metabolic activation and detoxication of carcinogenic N-nitrosodiethylamine in rat liver. J Agric Food Chem 2009, 57, 5071–5077, doi:10.1021/jf803973y.

37. Polk, A.; Vaage-Nilsen, M.; Vistisen, K.; Nielsen, D.L. Cardiotoxicity in cancer patients treated with 5-fluorouracil or capecitabine: a systematic review of incidence, manifestations and predisposing factors. Cancer treatment reviews 2013, 39, 974–984, doi:10.1016/j.ctrv.2013.03.005.

38. Sara, J.D.; Kaur, J.; Khodadadi, R.; Rehman, M.; Lobo, R.; Chakrabarti, S.; Herrmann, J.; Lerman, A.; Grothey, A. 5-fluorouracil and cardiotoxicity: a review. Ther Adv Med Oncol 2018, 10, 1758835918780140, doi:10.1177/1758835918780140.

39. Altena, R.; de Haas, E.C.; Nuver, J.; Brouwer, C.A.; van den Berg, M.P.; Smit, A.J.; Postma, A.; Sleijfer, D.T.; Gietema, J.A. Evaluation of sub-acute changes in cardiac function after cisplatin-based combination chemotherapy for testicular cancer. Br J Cancer 2009, 100, 1861–1866, doi:10.1038/sj.bjc.6605095.

40. Demkow, U.; Stelmaszczyk-Emmel, A. Cardiotoxicity of cisplatin-based chemotherapy in advanced non-small cell lung cancer patients. Respir Physiol Neurobiol 2013, 187, 64–67, doi:10.1016/j.resp.2013.03.013.

41. Dugbartey, G.J.; Peppone, L.J.; de Graaf, I.A. An integrative view of cisplatin-induced renal and cardiac toxicities: Molecular mechanisms, current treatment challenges and potential protective measures. Toxicology 2016, 371, 58–66, doi:10.1016/j.tox.2016.10.001.

42. El-Awady el, S.E.; Moustafa, Y.M.; Abo-Elmatty, D.M.; Radwan, A. Cisplatin-induced cardiotoxicity: Mechanisms and cardioprotective strategies. Eur J Pharmacol 2011, 650, 335–341, doi:10.1016/j.ejphar.2010.09.085.

43. Brown, D.A.; Perry, J.B.; Allen, M.E.; Sabbah, H.N.; Stauffer, B.L.; Shaikh, S.R.; Cleland, J.G.; Colucci, W.S.; Butler, J.; Voors, A.A., et al. Expert consensus document: Mitochondrial function as a therapeutic target in heart failure. Nat Rev Cardiol 2017, 14, 238–250, doi:10.1038/nrcardio.2016.203.

44. Varga, Z.V.; Ferdinandy, P.; Liaudet, L.; Pacher, P. Drug-induced mitochondrial dysfunction and cardiotoxicity. Am J Physiol Heart Circ Physiol 2015, 309, H1453–1467, doi:10.1152/ajpheart.00554.2015.

45. Gogvadze, V.; Orrenius, S.; Zhivotovsky, B. Mitochondria as targets for cancer chemotherapy. Semin Cancer Biol 2009, 19, 57–66, doi:10.1016/j.semcancer.2008.11.007.

46. Gorini, S.; De Angelis, A.; Berrino, L.; Malara, N.; Rosano, G.; Ferraro, E. Chemotherapeutic Drugs and Mitochondrial Dysfunction: Focus on Doxorubicin, Trastuzumab, and Sunitinib. Oxid Med Cell Longev 2018, 2018, 7582730, doi:10.1155/2018/7582730.

47. Beffagna, G. Zebrafish as a Smart Model to Understand Regeneration After Heart Injury: How Fish Could Help Humans. Front Cardiovasc Med 2019, 6, 107, doi:10.3389/fcvm.2019.00107.

48. Bournele, D.; Beis, D. Zebrafish models of cardiovascular disease. Heart Fail Rev 2016, 21, 803–813, doi:10.1007/s10741-016-9579-y.

49. Liu, J.; Stainier, D.Y. Zebrafish in the study of early cardiac development. Circ Res 2012, 110, 870–874, doi:10.1161/CIRCRESAHA.111.246504.

