## Supplementary material for "A polyphenol-rich extract of Olive Mill Wastewater Enhances cancer chemotherapy effects, while mitigating cardiac toxicity"

Suppl. Figure 1

| PHENOLIC COMPOUND | A009 L3 (g/L) | A009 L4 (g/L) |
|---|---|---|
| Hydroxytyrosol | 5.72 | 5.50 |
| Hydroxytyrosol glucoside | 1.69 | 1.91 |
| Verbascoside | 1.32 | 1.07 |
| 6’-p-coumaroyl secologanoside | 0.40 | 0.35 |
| b-hydroxyverbascoside isomer 1 | 0.14 | 0.23 |
| b-hydroxyverbascoside isomer 2 | 0.17 | 0.23 |
| Chlorogenic acid | 0.10 | 0.13 |
| Caffeoyl ester of secologanoside | 0.20 | 0.23 |
| Decarboxymethyloleuropein aglycon | 0.28 | 0.16 |
| Oleouropein aglycon | 0.22 | 0.21 |
| Tyrosol | - | 0.69 |
| Rutin | - | - |
| Luteolin-7-o-glucoside | - | - |

**L3**

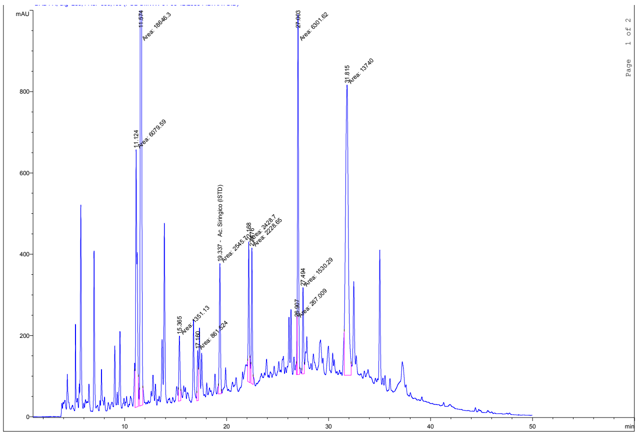


**L4**

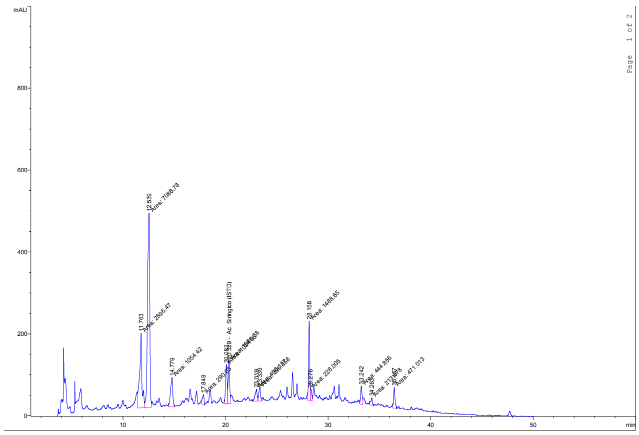

Suppl. Figure 2

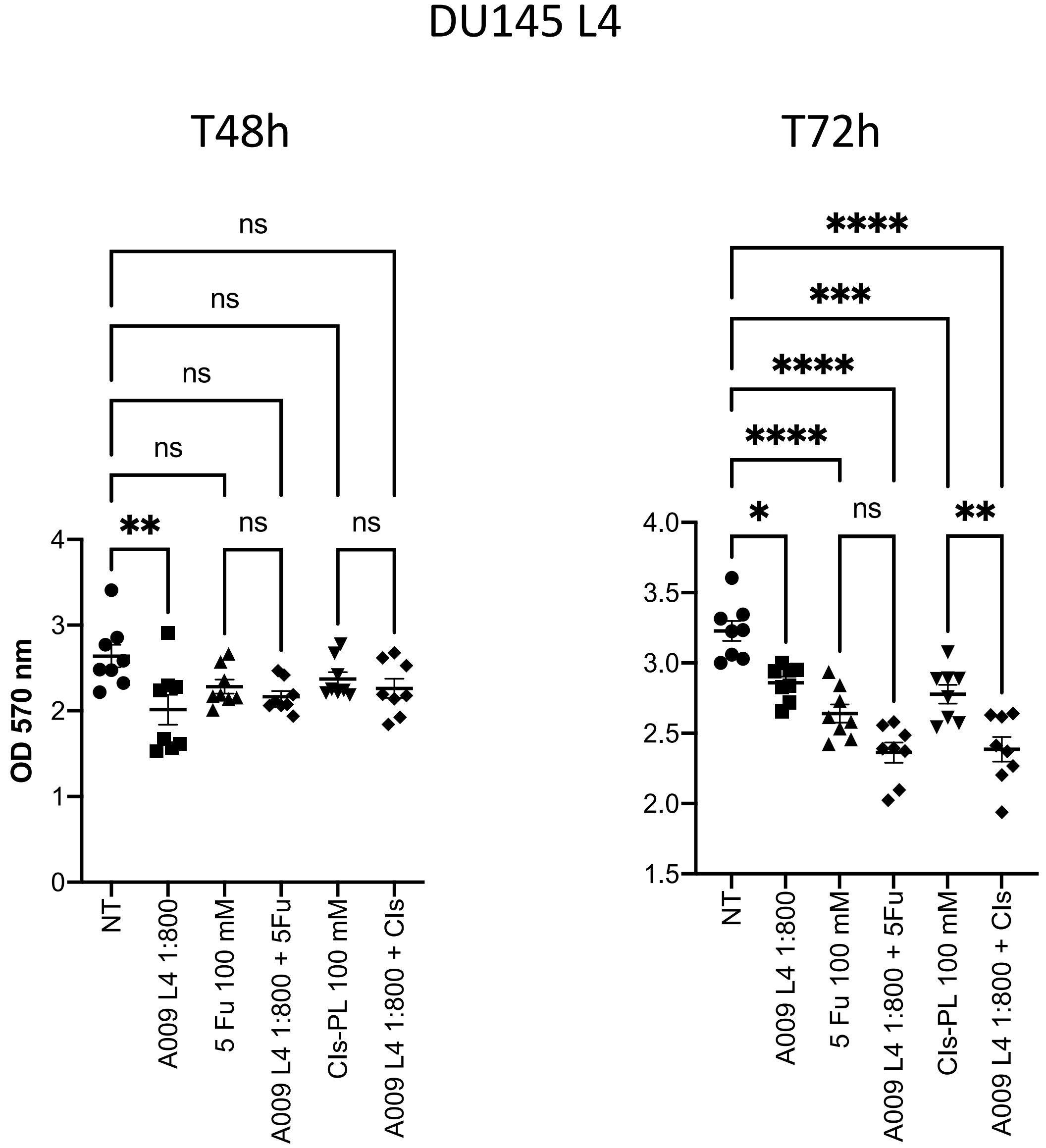
